# Ectopic expression of KRAS^G12D^ and p53^R167H^ in porcine mammary epithelial cells results in transformation and tumorigenesis

**DOI:** 10.1101/2021.06.20.449198

**Authors:** Neeley Remmers, Mark A. Carlson

## Abstract

We describe our initial studies in the development of an orthotopic, genetically-defined, large animal model of breast cancer, using immunocompetent pigs. Primary mammary epithelial cells were isolated from the porcine gland. Primary mammary cells were immortalized with hTERT, and then transformed cell lines were generated from these immortalized cells with oncogenic KRAS and dominant negative p53. The transformed cell lines outperformed the primary cells in terms proliferation, population doubling time, soft agar growth, 2D migration, and Matrigel invasion. Three transformed cell lines were selected based on *in vitro* performance, and were able to grow tumors when injected subcutaneously in nude mice, with undifferentiated morphology. Tumorigenic porcine mammary cell lines were generated in this report.

## INTRODUCTION

The annual incidence of breast cancer (BC) in women (all ages and races) in the United States increased from 0.102% in 1980 to a peak of 0.142% in 1999, and then decreased slightly, plateauing at ~0.131% from 2011-2017.^1^ As of 2017, a woman’s lifetime risk of developing BC in the U.S. is 12.9%.^1^ In 2021, the estimated number of new BC cases in the U.S. will be 281,550 (15.3% of all new cancer cases), with 43,600 estimated deaths (~20 per 100,000 in the general population, or 7% of all cancer deaths, or ~2% of all mortality in the U.S.).^1, 2^ All-stages 5-year survival for BC has improved from 75% in 1975 to 90% in 2016,^1^ secondary to earlier diagnosis and more efficacious therapy.^3^ However, survival with triple-negative breast cancer (TNBC; minimal/nil expression of the estrogen receptor, progesterone receptor, and epidermal growth factor receptor 2), which accounts for 10-20% of all BC,^4, 5^ is 10, 20, and 30% lower at stages 2, 3, and 4, respectively, compared to non-TNBC.^6^ So, there remain a need for improved management of BC, particularly TNBC.

Commonly utilized murine BC models^7–10^ include cell-line derived xenograft (CDX), patient derived xenograft (PDX), humanized CDX and PDX, and genetically engineered murine (GEM) model. CDX models are useful for BC genetics, but are poorly predictive of human response.^11^ PDX models may be more predictive for preclinical drug evaluation, but ultimately are limited by the host’s immunodeficient status.^10, 11^ This limitation has been somewhat countered by introducing components of the human immune system in the CDX and PDX models,^10, 11^ but graft *vs*. host issues continue to define the utility of the humanized models. GEM models^9, 12^ offer a wide range inducible, tissue-specific genetic alterations, are autochthonous, have immunocompetent hosts, and are genetically defined. BC can be modeled with any of the above murine systems.

Unfortunately, no murine model can overcome the limitation of inadequate subject size. Studies on ablation technologies, imaging modalities, tumor markers, chemotherapeutic pharmacokinetics, and other areas relevant to BC are difficult to perform in a subject whose mass is less than one-thousandth that of a typical patient. The availability of a large animal model of breast cancer, such as in the pig (which would approximate a human-sized subject), could address the size inadequacy of murine models. In the present report we describe our initial studies in deriving transformed, tumorigenic cell lines from primary cultures of porcine mammary epithelial cells (pMECs).

## METHODS AND MATERIALS

### Standards, rigor, reproducibility, and transparency

The animal studies of this report were designed, performed, and reported in accordance with both the ARRIVE recommendations (Animal Research: Reporting of *In Vivo* Experiments^13^) and the National Institutes of Health Principles and Guidelines for Reporting Preclinical Research.^14^

### Materials and animal subjects

All reagents were purchased through Thermo Fisher Scientific (www.thermofisher.com) unless otherwise noted. Athymic homozygous nude mice (Crl:NU(NCr)-*Foxn1^nu^*; female; 8-9 weeks old) were purchased from Charles River Laboratories, Inc. (www.criver.com). DNA sequencing was performed by the UNMC Genomics Core Facility (www.unmc.edu/vcr/cores/vcr-cores/genomics).

### Animal welfare

The animals utilized to generate data for this report were maintained and treated in accordance with the *Guide for the Care and Use of Laboratory Animals* (8^th^ ed.) from the National Research Council and the National Institutes of Health, and also in accordance with the Animal Welfare Act of the United States (U.S. Code 7, Sections 2131 – 2159). The animal protocols pertaining to this manuscript were approved by the Institutional Animal Care and Use Committee (IACUC) of the VA Nebraska-Western Iowa Health Care System (ID number 00998) and by the IACUC of the University of Nebraska Medical Center (ID number 16-133-11-FC). All procedures were performed in animal facilities approved by the Association for Assessment and Accreditation of Laboratory Animal Care International (AAALAC; www.aaalac.org) and by the Office of Laboratory Animal Welfare of the Public Health Service (grants.nih.gov/grants/olaw/olaw.htm). All surgical procedures were performed under isoflurane anesthesia, and all efforts were made to minimize suffering. Euthanasia was performed in accordance with the AVMA Guidelines for the Euthanasia of Animals.^15^

### Isolation of porcine mammary epithelial cells (pMECs)

Mammary tissue from a recently lactating domestic sow (age 3 years) that was scheduled for euthanasia for other reasons was obtained from a closed herd of research swine at the Eastern Nebraska Research and Extension Center (extension.unl.edu/statewide/enre) in Mead, NE. Immediately after sacrifice, the skin of a mammary mound was washed with chlorhexidine soap, and ~20 cm^3^ of mammary tissue was harvested with a scalpel. The tissue was placed on ice for ~1 h during transport to the laboratory. The tissue then was minced into ~1 mm cubes, and the fragments were enzymatically digested with 1 mg/mL of Collagenase D at 37°C for 1 h with gentle shaking. The digestate then was passed through a 70 μm nylon sieve, and cells were pelleted (600 *g* × 5 min) from the filtrate. The supernatant was discarded, the cell pellet was resuspended in whole media, which was defined as: DMEM (high glucose with L-glutamine; Thermo Fisher Scientific, cat. no. 12100-046) supplemented with 10% (final concentration) fetal bovine serum (FBS; Thermo Fisher Scientific, cat. no. 26140079) and 1% Antibiotic-Antimycotic Solution (Corning Inc., cat. no. 30-004-CI; cellgro.com).

Cell concentration in the resuspension was determined with a hemocytometer. This initial crude cell suspension then was were diluted and pipetted into a 96-well plate (1-10 cell/well, 100-200 μL/well). After 5-7 days of culture under standard conditions (defined as whole media, 37°C, 5% CO_2_), wells that contained cells with epithelial-like morphology were trypsinized and re-plated into a new 96-well plate, in order to dilute out any fibroblasts. Cells were passaged in this fashion at least four times, until no cells with fibroblast morphology were present. The resulting cells (primary pMECs; see Table tt02) were then seeded onto 6-well Corning plates, and then infected with human telomerase reverse transcriptase (hTERT)-containing lentivirus (AddGene) per the manufacturer’s instructions. The hTERT-LV treated pMECs (hpMECs) were expanded and used for subsequent transformation experiments.

**Table tt02.** Designations and definitions.

|  |  |
| --- | --- |
| pMECs | porcine mammary epithelial cells |
| hpMECs | pMECs infected with hTERT |
| LV-GKP | Lentivirus expressing <u>G</u>FP, <u>K</u>RAS<sup>G12D</sup>, and <u>p</u>53<sup>R167H</sup> |
| BGKP | hpMECs transformed with KRAS<sup>G12D</sup> and p53<sup>R167H</sup> (<u>B</u>reast <u>G</u>FP <u>K</u>RAS <u>p</u>53) |

### Generation of p53 and KRAS mutants and construction of expression vector

In order to generate the porcine p53^R167H^ mutant, wild-type p53 cDNA first was amplified from cervical lymph node tissue, which was obtained <5 min after euthanasia of a 4-month-old male domestic swine that had been on an unrelated research protocol. In brief, fresh nodal tissue was flash-frozen in liquid N_2_ and then pulverized with a mortar and pestle, with continual addition of liquid N^2^ during pulverization. The frozen powder then was placed into the first buffer solution of the QIAGEN RNEasy Mini Kit (cat. no. 74104; www.qiagen.com), and total RNA was isolated per the manufacturer’s instructions.

After isolation, the total RNA underwent reverse transcription to cDNA with a Verso cDNA Synthesis Kit (Thermo Fisher Scientific, cat. no. AB1453A), per the manufacturer’s instructions. The wild type p53 sequence was amplified out of the cDNA using the PCR primers shown in Table tt01, which flanked the p53 cDNA with *Sal*I and *Bam*HI restriction sites. Successful amplification of the wild-type p53 cDNA was confirmed by inserting the amplified candidate sequence into the TOPO^®^ vector (TOPO^®^ TA Cloning^®^ Kit; Invitrogen™/Life Technologies™, Thermo Fisher Scientific, cat. no. K202020) per the manufacturer’s instructions, followed by sequencing to confirm successful insertion.

**Table tt01.**
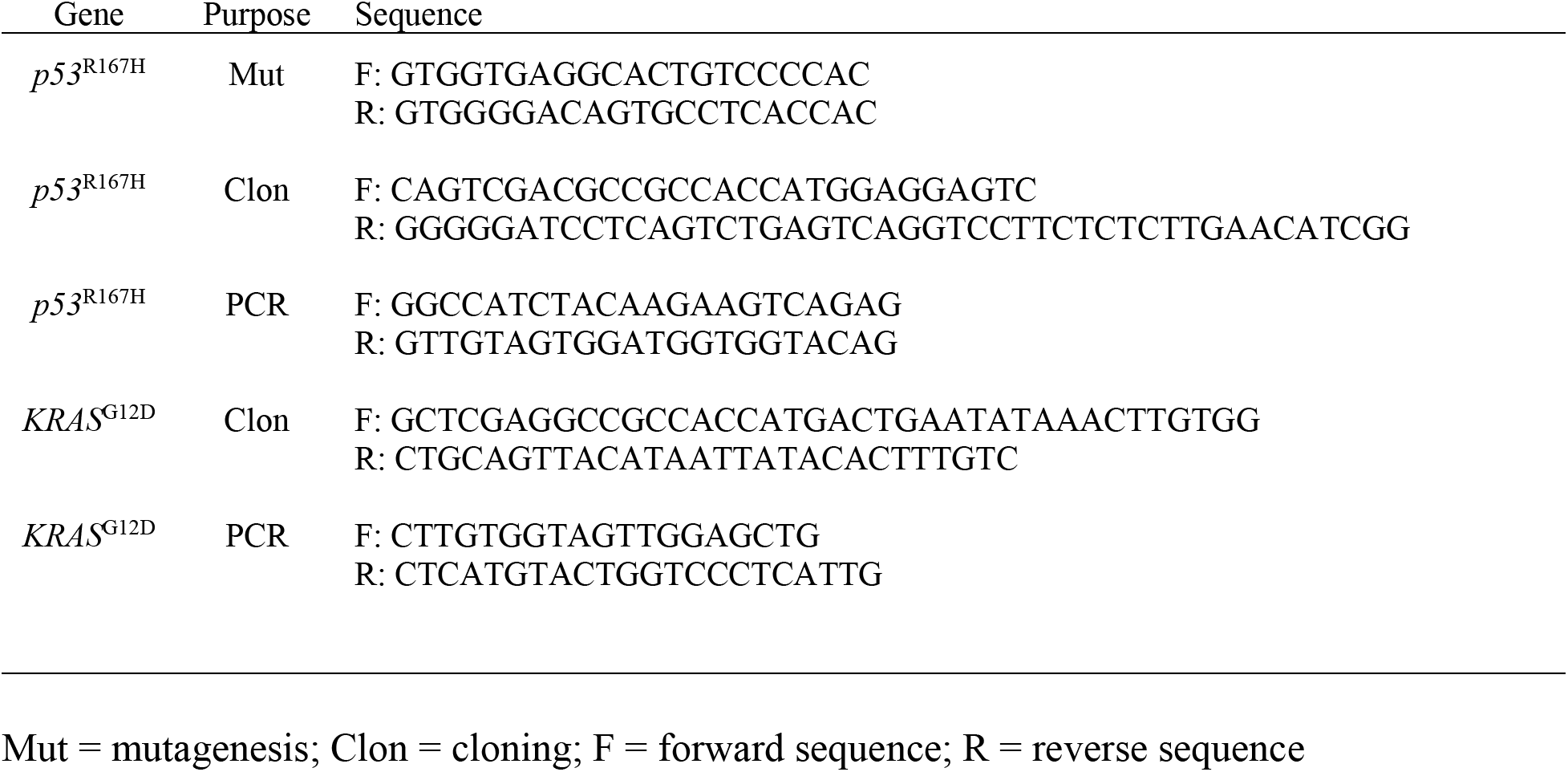
DNA sequences used for mutagenesis, cloning, PCR.

Site-directed mutation of wild-type p53 into p53^R167H^ was performed using Agilent Technologies’ QuickChange II Site-Directed Mutagenesis Kit (cat. no. 200523; www.genomics.agilent.com) with the mutagenic primers shown in Table tt01, per the manufacturer’s instructions. Presence of the p53^R167H^ mutation was verified by sequencing. The multiple cloning site of a pIRES2-AcGFP1 plasmid vector (Takara Bio USA, Inc., cat. no. 632435; www.clontech.com) was cut with *Sal*I and *Bam*HI, and the p53^R167H^ sequence then was ligated into this plasmid.

The source of the porcine KRAS^G12D^ mutant was the plasmid used to generate the p53/KRAS Oncopig.^16^ The KRAS^G12D^ cDNA was amplified out of this plasmid with primers (see Table tt01) that flanked the sequence with *Xho*I and *Pst*I restriction sites. The amplified product was inserted into the TOPO vector and verified by sequencing, as described above. The above pIRES2-AcGFP1 plasmid (already containing the p53^R167H^ sequence) then was cut with *Xho*I and *Pst*I, and the KRAS^G12D^ sequence was ligated into this plasmid, producing a pIRES2-AcGFP1 plasmid which contained both mutant cDNAs (KRAS^G12D^ and p53^R167H^) within its multiple cloning site (KRAS^G12D^ upstream); see Fig. tf02.

**Fig. tf01.**
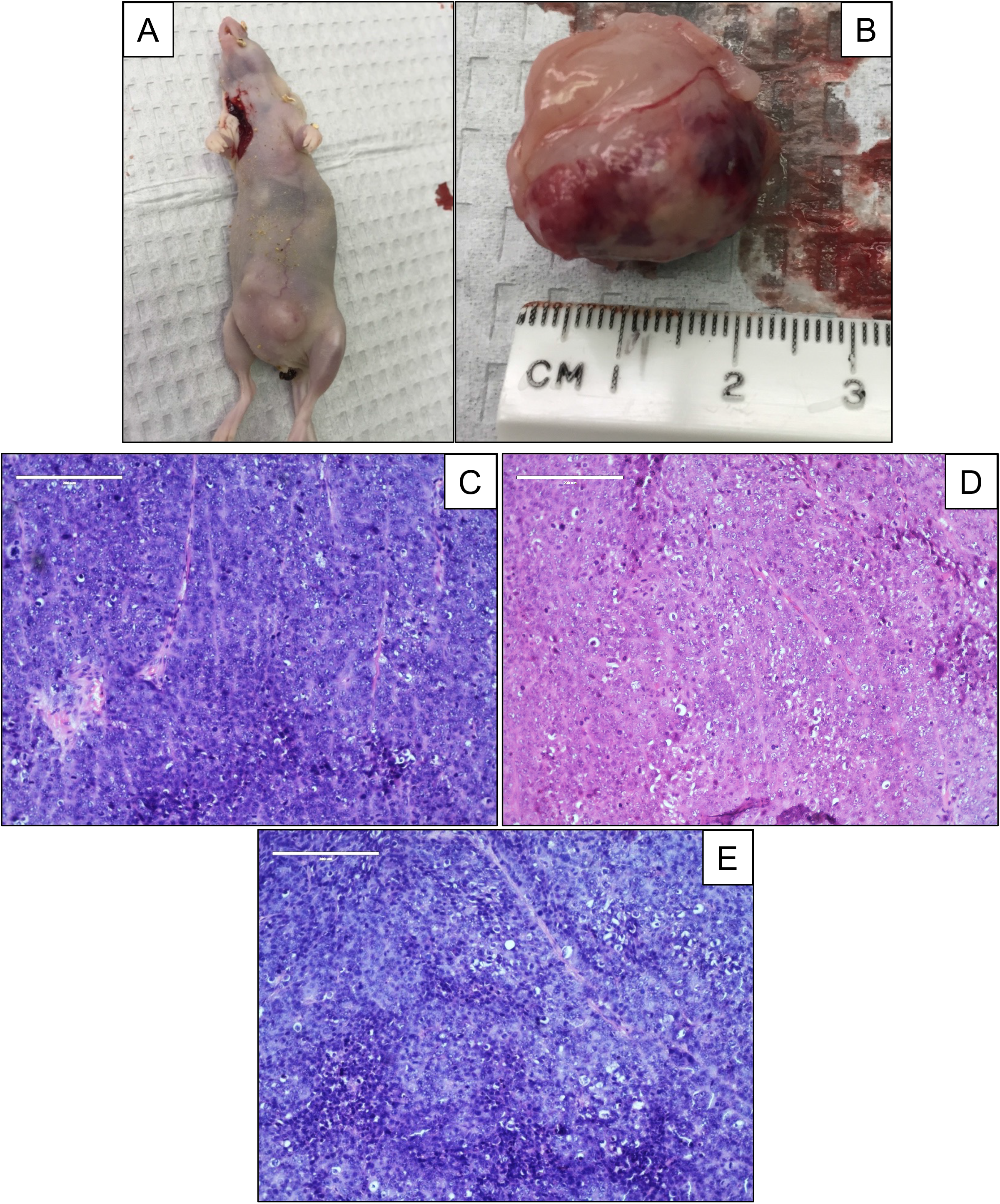
Implantation of transformed pMECs into nude mice. Mice were injected with 1M cells/injection (4 injections/mouse) along the nipple lines, and followed for up to 10 weeks. (A) Typical implanted mouse (BGKP 9.1 line), euthanized at 6 weeks. (B) Explanted xenograft tumor from panel A. (C-E) H&E histology of xenograft tumors explanted from nude mice (BGKP cell lines): (C) 9.1; (D) 9.4; (E) 16.5. Bar = 200 μm.

**Fig. tf02.**
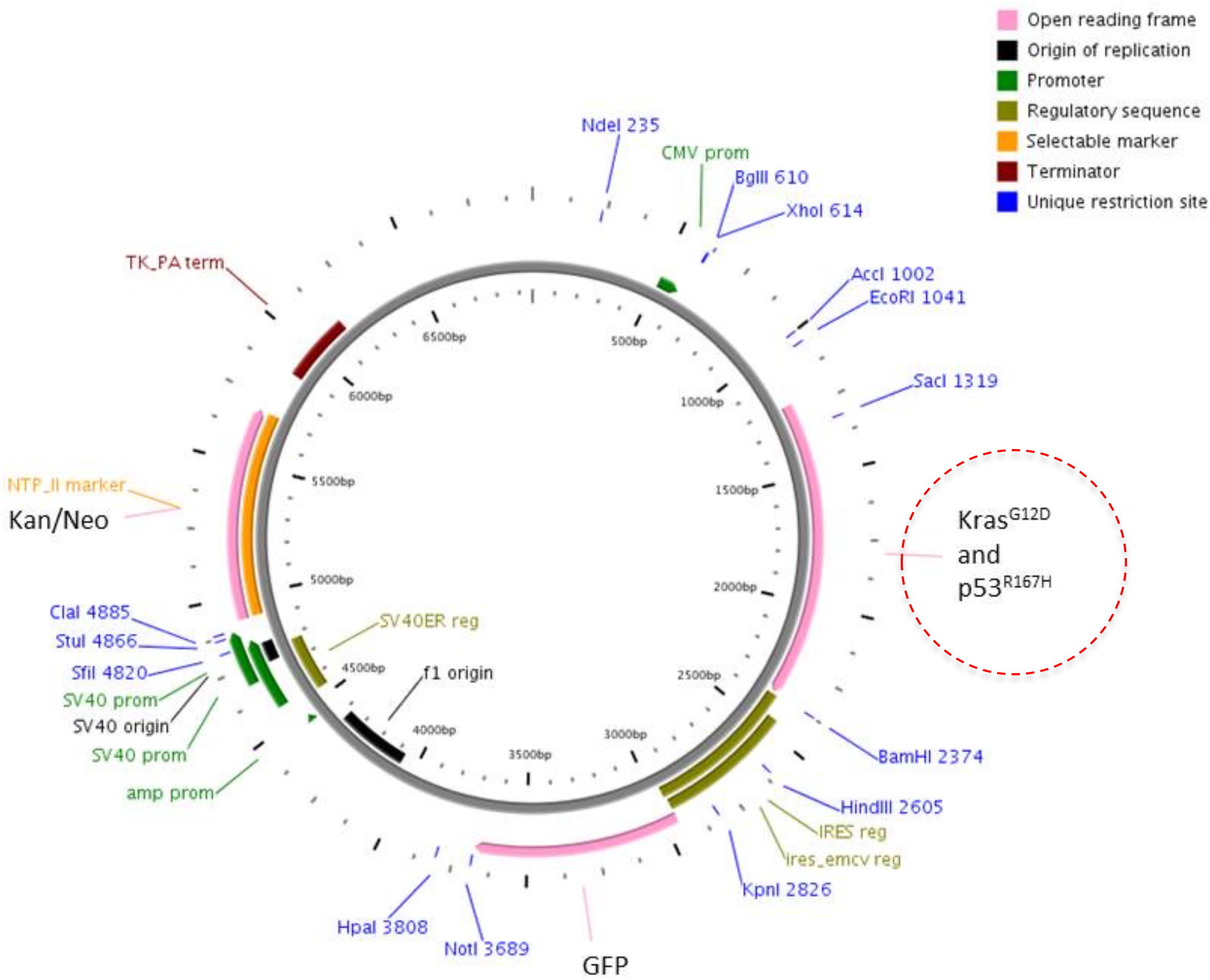
Vector Information. Plasmid used for construction of the lentiviral vector subsequently utilized for introduction of KRAS^G12D^ and p53^R167H^ (dashed red circle) into pMECs. Expression of the mutant genes is under control of the CMV promoter. Construct also expresses GFP via an IRES construct.

This newly-constructed plasmid (Fig. tf02), hereafter designated as GKP (G = AcGFP1; K = KRAS^G12D^; P = p53^R167H^), was transformed into One Shot™ Stbl3™ Chemically Competent *E. coli* (Invitrogen™/Thermo Fisher Scientific, cat. no. C737303), per the manufacturer’s instructions, and plasmid DNA subsequently was isolated using a QIAGEN Plasmid Maxi Kit (cat. no. 12162), per the manufacturer’s instructions. This plasmid then was utilized to construct a lentiviral vector, which ultimately was produced with Takara’s Lenti-X™ 293T cells (Clontech, cat. no. 632180) per the manufacturer’s instructions, to generate infectious lentiviral particles that would direct expression of AcGFP1, KRAS^G12D^, and p53^R167H^ mutants in transduced cells.

### Cell transformations

hpMECs were grown in T75 flasks to 80% confluency under standard conditions. The media then was exchanged with 2-3 mL of supernatant from non-lysed Lenti-X™ 293T cells (containing GKP viral particles) with 2 μg/mL polybrene (cat. no. TR1003, Thermo Fisher Scientific). After 24-48 h at 37°C, LV-infected hpMECs were re-seeded into 6-well plates under standard conditions and grown to 80% confluency. An exchange with whole media containing 2 μg/mL G418 aminoglycoside antibiotic then was performed; the G418 dose was chosen based on preliminary dose-response studies against non-treated epithelial cells. After 24 h, a whole media exchange was done, and the presence of transduced cells was determined with inverted GFP fluorescent microscopy of living cells. Cells expressing GFP after this process (i.e., the putative KRAS/p53 transformed cells) were designated as BGKP lines (see Table tt02), of which there were two series, “9” and “16”.

### PCR

Cell and tissue RNA was isolated using the QIAGEN RNEasy Mini Kit. Purified RNA then was used to generate cDNA using the Verso cDNA Synthesis Kit. The Platinum^®^ Blue PCR Supermix (Invitrogen™/Life Technologies, cat. no. 12580) subsequently was used for all PCR reactions. Amplified products were separated with agarose gel electrophoresis, and then visualized using a UV-light box. qPCR was performed using the PowerUp™ SYBR^®^ Green Master Mix (Applied Biosystems™/ Thermo Fisher Scientific, cat. no. A25741) per manufacturer’s protocol, and run on an Applied Biosystems™ 7500 Fast Dx Real-Time PCR Instrument. Fold changes in gene expression were calculated using the comparative *C*_T_ method.^17^ All primers used are listed in Table tt01.

### Soft agar assay

A standard soft agar assay^18^ was used to determine anchorage independent growth. A base layer of 1% agarose was plated into 6-well plates. A total of 2,500 cells/well were mixed with 0.7% agarose and plated on top of the base layer. The plates were incubated under standard conditions for 21 days. The cells then were stained with crystal violet, and counted using an inverted microscope. Cells were plated in triplicate, and total counts from all three wells were averaged.

### Migration assay

A standard scratch assay (monolayer wounding)^19^ was performed to determine cellular migration rate. Cells were plated in triplicate into 6-well plates. A horizontal scratch using a 10 μL pipet tip was made in each well. After washing away scratched-off cells, baseline images along the scratch were obtained, the plates were incubated under standard conditions, and subsequent images were captured at 3, 6, 9, 12, and 15 h after the initial scratch. ImageJ software (imagej.nih.gov/ij) was used to measure the distance between the two migrating cellular fronts (scratch edges) at 3-5 locations along the scratch. Average distance at each time point was plotted to generate the migratory rate (μm/h).

### Invasion assay

BioCoat™ Matrigel™ Invasion Chambers (Corning™, Thermo Fisher Scientific, cat. no. 08-774) were plated with 50,000 cells (upper chamber) in triplicate, and incubated under standard conditions for 24 h. The media from the upper chamber then was removed, and any cells remaining in the upper chamber were removed using a cotton swab. Cells that had migrated to the bottom of the membrane were stained using a Kwik-Diff™ kit (Shandon™, Thermo Fisher Scientific, cat. no. 9990701). Membranes were mounted onto glass slides, and cells per high-power field were counted using ImageJ software.

### Population doubling assay

Cells were plated in 6-well plates (10,000 cell/well), and cultured under standard conditions. Triplicate plates then were trypsinized on days 1, 2, 3, 4, 6, and 8, and cells were counted with a hemocytometer. Cell number *vs*. day was plotted to determine the day range in which linear growth was achieved. The data from this linear growth phase were used to determine population doubling time (DT) using the formula: DT = (Δt) × ln(2) ÷ ln(N_*f*_/N_*i*_) where Δt = time interval between initial and final cell count, N_*f*_ = cell count at final time, and N_*i*_= cell count at initial time.

### Proliferation assay

Relative cell proliferation rates were assayed using an MTT (3-(4,5-dimethylthiazol-2-yl)-2,5-diphenyltetrazolium bromide) assay kit (Vybrant™ MTT Cell Proliferation Assay Kit, Invitrogen™, Thermo Fisher Scientific, cat. no. V13154). Cells were plated in triplicate in a 96-well plate (5,000 cell/well), and cultured under standard conditions for 48 h. MTT reagent then was added to the cells per the manufacturer’s instructions, followed by addition of the solvent solution 3.5 h later. Absorbance was measured with a plate reader 3.5 h after solvent addition. Mean absorbance was normalized to absorbance from hpMECs to calculate fold-difference in proliferation.

### Subcutaneous implantation into nude mice

Subcutaneous implantation of BGKP cell lines into immunodeficient (athymic) mice was performed as previously described,^20^ with some modifications. BGKP cells (along with control hpMECs) were trypsinized, counted, and resuspended in DMEM at a concentration of 1 × 10^7^ viable cells/mL. Nude mice (50% female; maintained in microisolator cages with soft bedding and fed regular chow) were randomized into treatment groups using an online randomization tool (www.randomizer.org). Mice then were injected with 1 × 10^6^ cells (100 μL) per injection, 1 injection per mouse, placed ventrally along the nipple line, under brief isoflurane inhalational anesthesia, administered with a Matrx VMS^®^ small animal anesthesia machine, within a small animal operating room. Subjects were observed for 10 weeks or until tumors reached 1.5 cm in diameter by physical exam, and then subjects were euthanized with CO_2_ asphyxiation. At necropsy all gross tumor was measured and collected, portions underwent formalin fixation and paraffin embedding, and sections subsequently underwent H&E or immunohistochemical staining as described above. An independent, a blinded pathologist analyzed the stained sections to determine whether tumors were epithelial in origin, and if they displayed malignant features.

### Statistics and power analysis

Data are reported as mean ± standard deviation. Groups of continuous data were compared with ANOVA and the unpaired t-test. Categorical data were compared with the Fisher or Chi square test. For the power analysis of the murine subcutaneous tumor implant assay, tumor diameter was selected as the endpoint. Setting alpha = 0.05 and power = 0.8, ten mice per group were needed across three groups to detect a difference in means of 30% with the standard deviation estimated at 20% of the mean.

## RESULTS

### Porcine mammary gland

Examination of breast tissue in a nulliparous female pig was performed to illustrate relevant gross and microscopic anatomy. Evidence of mammary bud development is evident as early as 4 mo of age in domestic female swine (Fig. tf05A). The nipple complex in the pig contains two main ductal orifices (Fig. tf05G). These orifices can be cannulated with the unaided eye using a small diameter (30 gauge) needle (Fig. tf05H), though loupe magnification can assist with this procedure. If the duct of an explanted nipple is injected with India ink after needle cannulation, ink can be seen emerging from multiple smaller ducts in the subcutaneous tissue, confirming intraductal placement of the needle. (Fig. tf05I). A histologic sagittal H&E section of the nipple confirms the presence of a main duct (Fig. tf05C-D), along with the presence of primitive structures resembling terminal ductal lobular units (TDLU; Fig. tf05D-E). In the pig, a lobule is lined with two layer of epithelium: an outer myoepithelial layer, and a luminal epithelial layer (Fig. tf05F), all contained within a basement membrane.

### Isolation and transformation of primary pMECs

pMECs cultured and passaged from minced porcine mammary tissue displayed epithelial morphology under phase microscopy (Fig. tf03A) and stained for CK5 (Fig. tf03B). Based on these results, we concluded that we had primary pMECs that could be immortalized with hTERT. Infection of the subsequent lines of the hpMECs with LV-GKP along with antibiotic selection yielded eight lines of BGKP cells (Table tt02). Typical transformed morphology and GFP fluorescence is shown in Fig. tf03C-D, respectively. These cells lines all over-expressed KRAS and p53 by immunoblot (Fig. tf03E), and had elevated levels of KRAS and p53 transcripts by qPCR (Fig. tf03F-G). Based on these data, we concluded that we had successfully transduced the hpMEC cells with the KRAS and p53 mutants.

**Fig. tf03.**
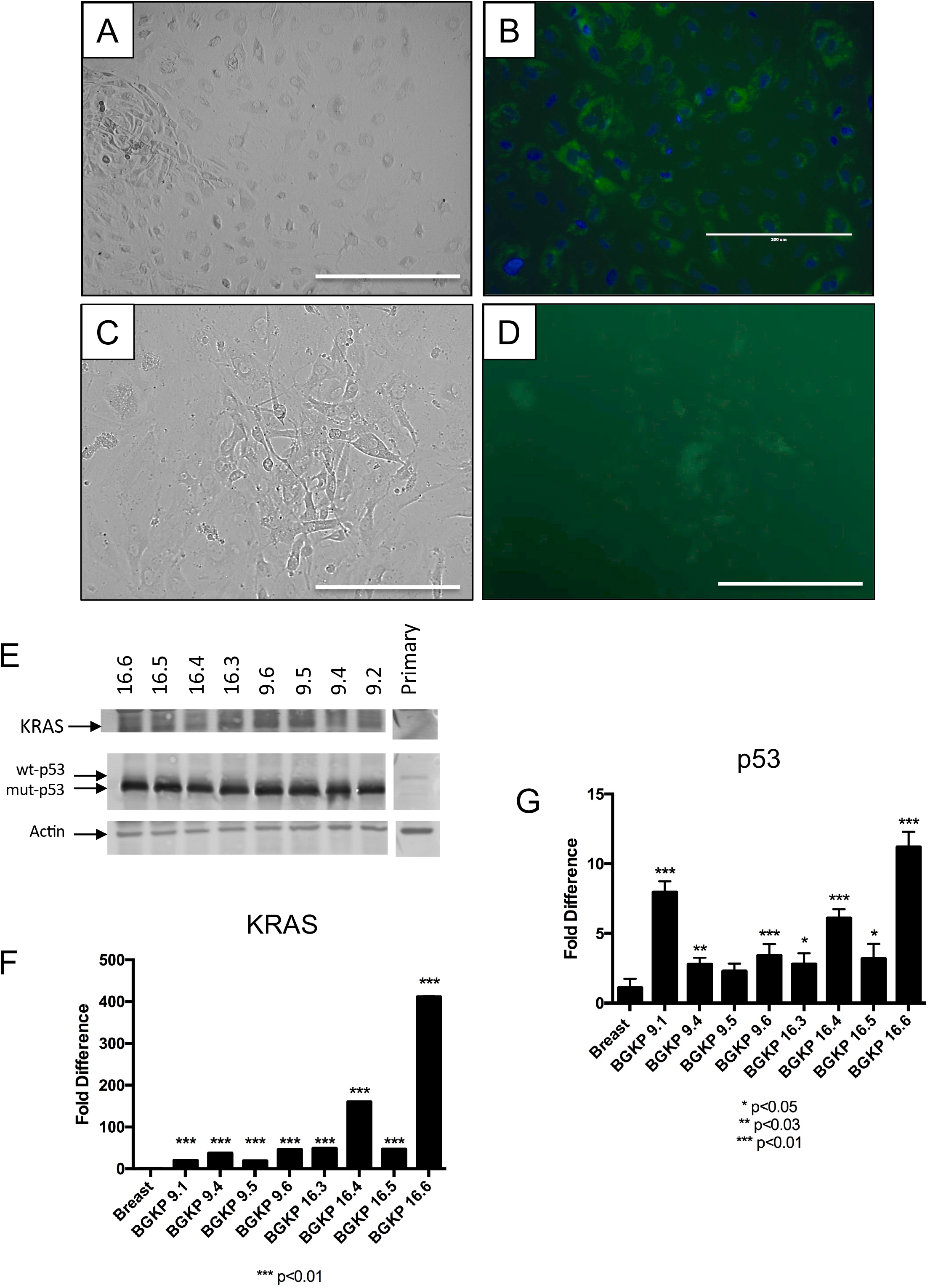
LV insertion of KRAS and p53 mutants into pMECs. (A) Primary pMECs, phase image (bar = 400 μm). (B) Primary pMECs, cytokeratin 5 immunohistochemistry (bar = 200 μm). (C) hpMEC after infection with LV-GKP, phase (bar = 200 μm). (D) Panel C, GFP fluorescence. (E) Immunoblot of KRAS^G12D^ and p53 in lysates of BGKP cell lines *vs*. primary pMECs (“primary”). (F-G) qPCR of KRAS and p53 in BGKP cell lines *vs*. hpMECs (“breast”). Results of unpaired t-testing shown (BGKP line *vs*. hpMEC).

### *In vitro* assays of transformation

We next wanted to characterize the newly derived BGKP cell lines using *in vitro* assays of transformation. In a monolayer wound healing (scratch) assay (Fig. tf04), all BGKP lines migrated over the scratch wound at least twice as fast as the hpMECs (Fig. tf04E). In a cell-count based assay of population growth performed over 8 days (Fig. tf07), 7 of the 8 BGKP lines had faster population growth than the hpMECs; one line (BGKP 16.3) was slower than the hpMECs. In a related dye-based proliferation assay performed over 48 h (Fig. tf06A), all 8 BGKP lines demonstrated higher proliferative capacity compared to hpMECs. In a colony-forming soft agar assay (Fig. tf06B), 4 BGKP lines had markedly higher colony formation compared to hpMECs, while two others had marginally higher colony formation. In a Matrigel^®^ invasion assay (Fig. tf06C), 3 of the 8 BGKP lines (9.1, 9.4, and 16.5) had greater invasive ability compared to the hpMECs. We concluded from these data that at least some of our BGKP lines (particularly 9.1, 9.4, and 16.5) demonstrated convincing (based on multiple assays) *in vitro* behavior that was consistent with transformation.

**Fig. tf04.**
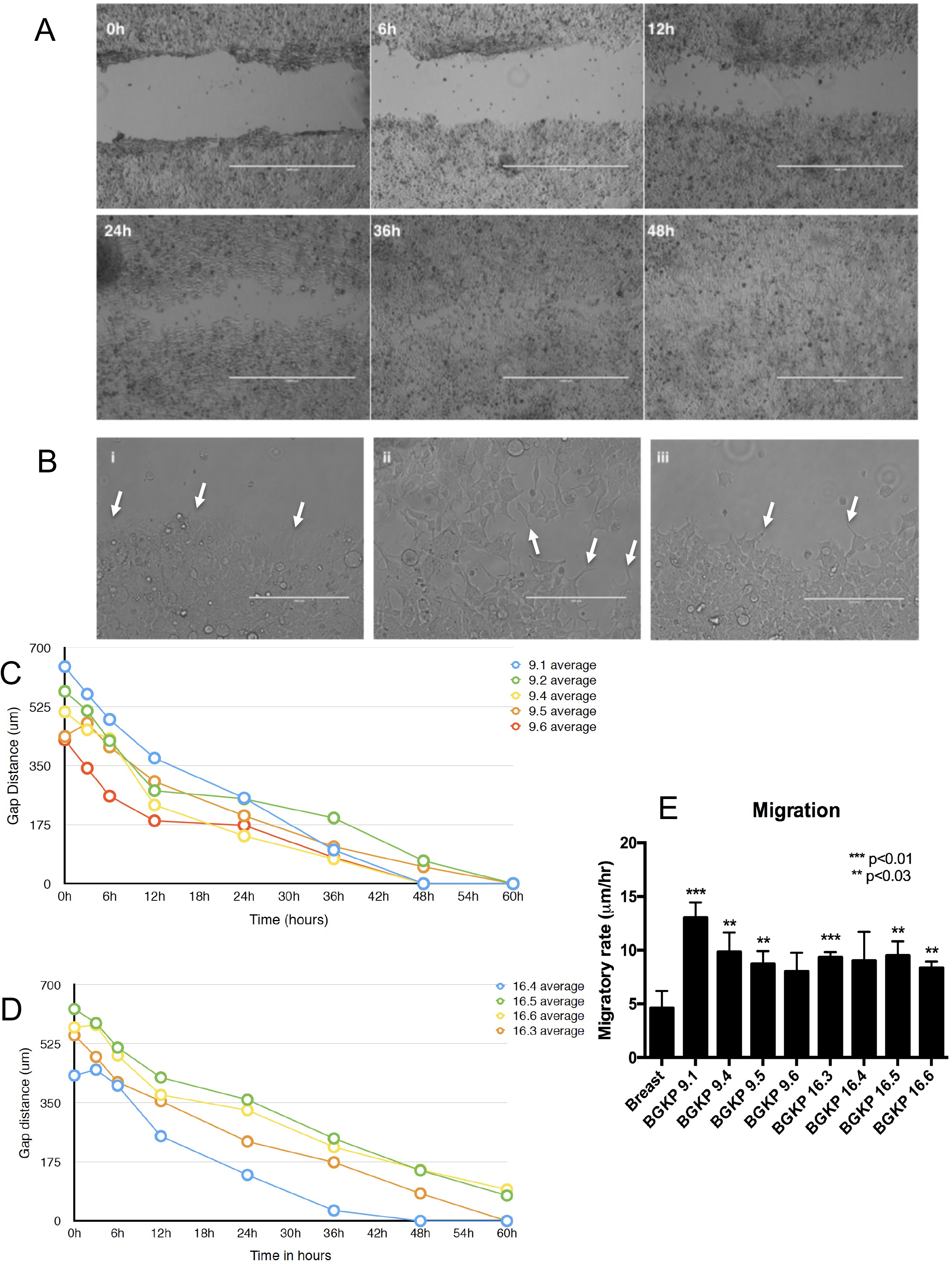
*In vitro* wound healing (scratch) assay. (A) Phase images 0 to 48 h after a single scratch made to cell line BGKP 9.4; bar = 400 μm. (B-i) Lamellipodia developing on border of BGKP 9.4 cells 3 h after scratching (arrows); bar = 100 μm. (B-ii) Migratory cells within the area of the BGKP 9.4 scratch at 24 h; (B-iii) Migratory border of cell line BGKP 16.5. (C-D) Plots of gap distance in the scratch assay for BGKP cell line series 9 and 16, respectively. Each point represents the mean of 3 gap distances (standard deviations were smaller than the data points). (E) Cell front migratory rates derived from the gap distance data for the BGKP cell lines *vs*. hpMECs (“Breast”). Results of unpaired t-testing shown (BGKP line vs. hpMECs).

**Fig. tf05.**
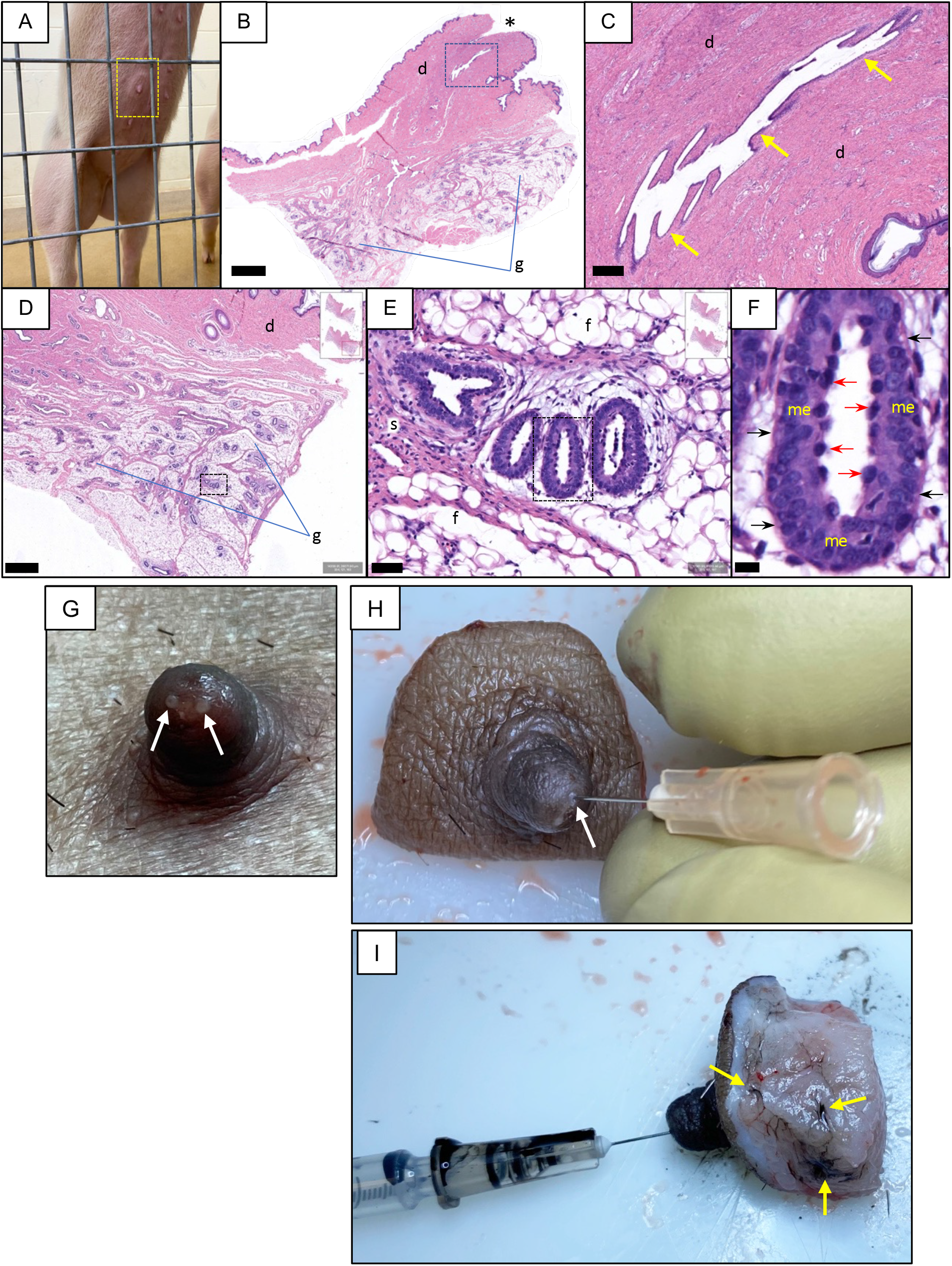
Porcine mammary gland. (A) Nulliparous domestic female pig, 4 mo/41 kg, ventral aspect, on hind legs, with visible breast mounds (dashed box). (B) Nipple from a nulliparous female pig; sagittal H&E histology; asterisk = nipple orifice; d = dermis; g = glandular tissue; bar = 2 mm. (C) Zoom of box in panel B, showing one of the main lactiferous ducts (yellow arrows); bar = 200 μm. (D) Glandular tissue in the same subject; bar = 800 μm. (E) Box in panel D, showing a terminal ductal lobular unit (TDLU); s = stroma; f = fat; bar = 50 μm. (F) Box in panel E; red arrows = inner (luminal) epithelium; black arrows = basement membrane; me = outer myoepithelium; bar = 10 μm.(G) Close-up of nipple *in vivo*, demonstrating two mammary ductal orifices (arrows). Diameter of nipple is 6-7 mm. (H) Nipple complex *ex vivo*. A ductal orifice (arrow) has been cannulated with a 30g needle. (I) Injection of ductal orifice with 50 μL India ink. Note that some ink has emerged through ducts within subcutaneous region of nipple complex (arrows), confirming intraductal injection.

**Fig. tf06.**
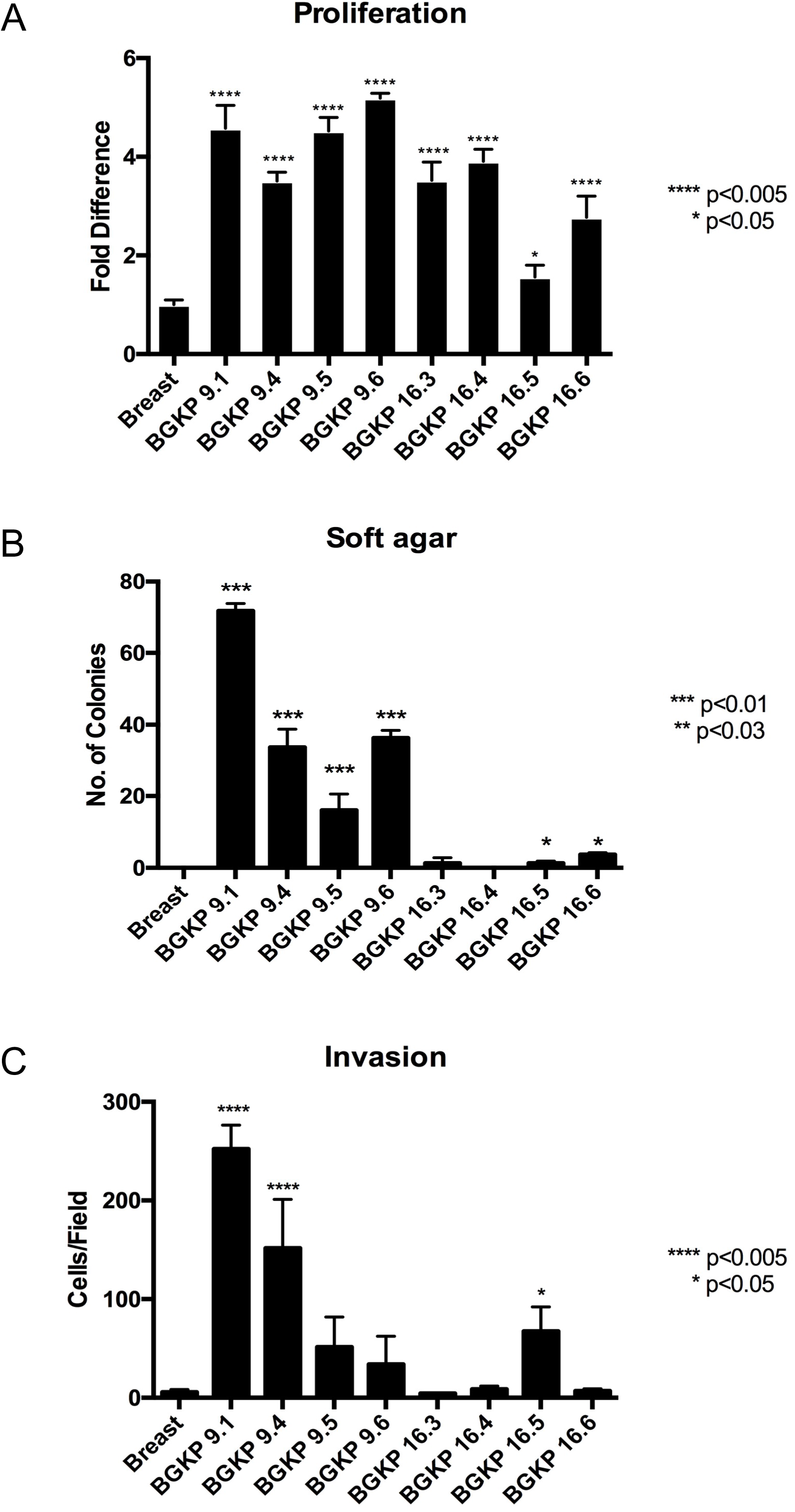
*In vitro* transformation assays of BGKP cell lines. (A) Cellular proliferation. (B) Soft agar colony growth. (C) Matrigel^®^ invasion. Breast = hpMECs; results of unpaired t-testing shown (BGKP cell line *vs*. hpMECs).

**Fig. tf07.**
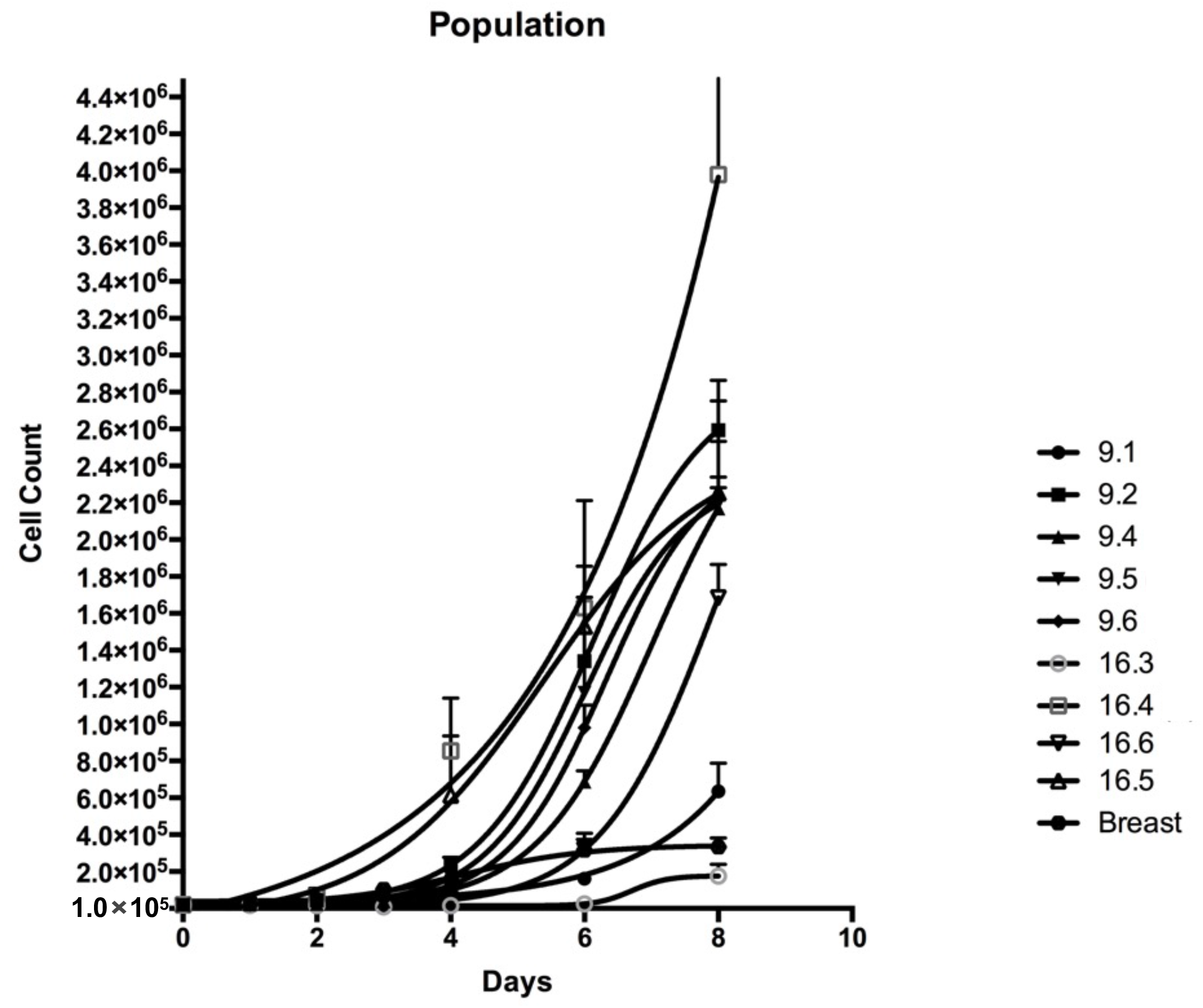
*In vitro* population growth of BGKP cell lines. Each data point represents the mean of 3 individual plates; error bars = standard deviations; Breast = primary pMECs.

### Subcutaneous implantation of BGKP lines into immunodeficient mice

In order to determine if the BGKP cell lines were tumorigenic properties *in vivo*, we utilized subcutaneous cell implantation into homozygous athymic mice. Based on the above *in vitro* data, we selected three BGKP lines (9.1, 9.4, and 16.5) for implantation into nude mice (Fig. tf01); see results in Table tt03. Control injections with non-transformed hpMECs did not produce any tumors. Histological analysis of these subcutaneous xenografts revealed mucin-producing, well vascularized tumors with large areas of necrosis. The tumors were pleomorphic and undifferentiated with little to no desmoplasia. All three BGKP tumors stained for E-cadherin, but poorly for EpCAM (data not shown).

**Table tt03.** Tumor xenografts.

| BGKP line | Tumors | *Mean diameter (cm) |
| --- | --- | --- |
| 9.1 | 2/8 | 1.1 |
| 9.4 | 1/8 | 0.7 |
| 16.5 | 5/8 | 1.0 |

8 mice/line; single injection of 1M cells

## DISCUSSION

Other groups have transformed porcine mammary epithelial cells *in vitro* with SV40 large T antigen insertion^21^ or BRCA1 knockdown,^22^ with evidence of tumorigenicity in immunodeficient mice.^21^ In addition, a *BRCA1* haploinsufficient Yucatan minipig was generated with somatic cell nuclear transfer,^23^ but postnatal survival of cloned piglets was ≤18 days for reasons that were unclear (*BRCA1*+/– mice are phenotypically normal^24^). To be clear, at present there is no porcine model of BC. Of note, while the KRAS/p53 Oncopig^16^ can generate subcutaneous tumors if injected with AdCre, this transgenic pig is not a BC model, but rather a KRAS-driven nonspecific tumor generator. Cellular transformation is nonspecific in the Oncopig; multiple cell types can be transformed, not just epithelial cells.

Our goal with the present line of experiments was to generate tumorigenic mammary cell lines that could eventually be used in an immunocompetent porcine model of BC. Current murine models of BC will continue to be helpful, particularly for the study of molecular mechanisms. However, murine models are limited in their ability to replicate human biology and size, so a large animal model of BC likely would enhance our ability to develop and test new diagnostic and treatment modalities for this disease. The data presented herein demonstrated that wild type porcine mammary epithelial cells can be transformed with modulation of common tumor-associated target genes, and that these transformed cells subsequently can grow tumors in immunodeficient mice. These data may provide a pathway for the construction of an orthotopic porcine model of BC, namely, implantation of tumorigenic hpMECs into the porcine mammary gland. Ultimately we would like to produce an orthotopic, genetically-defined, porcine model of breast cancer, using immunocompetent pigs.

In our initial derivation of tumorigenic mammary cell lines for the pig, we selected KRAS and p53 mutants to transform our hTERT-immortalized pMECs. The rationale for the initial selection of these two mutants to transform porcine mammary cells was based on the availability of the mutant KRAS^G12D^ sequence from our work on a porcine model of pancreatic cancer, and the knowledge that these two sequences together could transform cultures of porcine pancreatic ductal epithelial cells and produce pancreatic cancer in the Oncopig.^25–27^ While the *KRAS*^G12D^ mutation occurs in BC, it is less common than other mutations involving *BRCA1* or *PIK3CA*.^28–30^ Our future work in porcine BC modeling will focus on utilizing genetic edits of *BRCA1*, *PIK3CA*, and *TP53*.

## ACKNOWLEDGEMENTS

This study is the result of work supported in part with resources and the use of facilities at the Omaha VA Medical Center (Nebraska-Western Iowa Health Care System). The work was supported by seed grants from the University of Nebraska Medical Center and the Eppley Cancer Center (now Buffett Cancer Center) of Omaha, Nebraska. The authors would like to acknowledge the technical assistance of Gerri Siford and Chris Hansen. The authors also would like to thank Dr. Lawrence B. Schook and Dr. Laurie A. Rund at the University of Illinois for the gift of their plasmid that contained the KRAS^G12D^ mutation, along with comments, insights and suggestions for its use in this project.

## Notes

Conflicts of Interest: The authors declare no relevant conflicts of interest.

### Competing Interest Statement

The authors have declared no competing interest.

